# Improved annotation of the insect vector of Citrus greening disease: Biocuration by a diverse genomics community

**DOI:** 10.1101/099168

**Authors:** Surya Saha, Prashant S Hosmani, Krystal Villalobos-Ayala, Sherry Miller, Teresa Shippy, Mirella Flores, Andrew Rosendale, Chris Cordola, Tracey Bell, Hannah Mann, Gabe DeAvila, Daniel DeAvila, Zachary Moore, Kyle Buller, Kathryn Ciolkevich, Samantha Nandyal, Robert Mahoney, Joshua Van Voorhis, Megan Dunlevy, David Farrow, David Hunter, Taylar Morgan, Kayla Shore, Victoria Guzman, Allison Izsak, Danielle E Dixon, Andrew Cridge, Liliana Cano, Xialong Cao, Haobo Jiang, Nan Leng, Shannon Johnson, Brandi L Cantarel, Stephen Richardson, Adam English, Robert G Shatters, Chris Childers, Mei-Ju Chen, Wayne Hunter, Michelle Cilia, Lukas A Mueller, Monica Munoz-Torres, David Nelson, Monica F Poelchau, Joshua B Benoit, Helen Wiersma-Koch, Tom D’elia, Susan J Brown

**Affiliations:** Boyce Thompson Institute, Ithaca, NY 14853, USA; Indian River State College, Fort Pierce, FL 34981, USA; Kansas State University, Division of Biology, Manhattan, KS 66506, USA; University of Cincinnati, Cincinnati, OH 45220, USA; Cornell University, Ithaca, NY 14853, USA; University of Puget Sound, Tacoma, WA 98416, USA; University of Otago, North Dunedin, Dunedin 9016, New Zealand; University of Florida/ IFAS Indian River Research and Education Center, Plant Pathology, Ft. Pierce, FL 34945, USA; Los Alamos National Laboratory, NM 87544, USA; UT Southwestern Medical Center, Bioinformatics Core Facility, Department of Bioinformatics, Dallas, TX 75390, USA; Baylor College of Medicine, i5K Arthropod Genomics, Houston, TX 77030, USA; Baylor College of Medicine, Human Genome Sequencing Center, Houston, TX 77030, USA; Illumina Inc., San Diego, CA 92122, USA; Oklahoma State University, Department of Biochemistry and Molecular Biology, Stillwater, OK 74074, USA; Oklahoma State University, Department of Entomology and Plant Pathology, Stillwater, OK 74074, USA; USDA Agricultural Research Service, National Agricultural Library, Beltsville, MD 20705, USA; National Taiwan University, Graduate Institute of Biomedical Electronics and Bioinformatics, Taipei 10617, TW; USDA ARS, U. S. Horticultural Research Laboratory, Ft. Pierce, FL 34945, USA; USDA ARS, Emerging Pests and Pathogens Research Unit, Ithaca, NY 14853, USA; Cornell University, Plant Pathology and Plant-Microbe Biology Section, School of Integrative Plant Science, Ithaca, NY 14853, USA; Cornell University, Plant Breeding and Genetics Section, School of Integrative Plant Science, Ithaca, NY 14853, USA; Lawrence Berkeley National Laboratory, Environmental Genomics and Systems Biology, Berkeley, CA 94720, USA; The University of Tennessee Health Science Center, Department of Microbiology, Immunology and Biochemistry, Memphis, TN 38163, USA

**Keywords:** Asian citrus psyllid, biocuration, i5k workspace at NAL, training, annotation, insect immunity

## Abstract

The Asian citrus psyllid (*Diaphorina citri* Kuwayama) is the insect vector of the bacterium *Candidatus* Liberibacter asiaticus (CLas), the pathogen associated with citrus Huanglongbing (HLB, citrus greening). HLB threatens citrus production worldwide. Suppression or reduction of the insect vector using chemical insecticides has been the primary method to inhibit the spread of citrus greening disease. Accurate structural and functional annotation of the Asian citrus psyllid genome, as well as a clear understanding of the interactions between the insect and CLas, are required for development of new molecular-based HLB control methods. A draft assembly of the *D. citri* genome has been generated and annotated with automated pipelines. However, knowledge transfer from well-curated reference genomes such as that of *Drosophila melanogaster* to newly sequenced ones is challenging due to the complexity and diversity of insect genomes. To identify and improve gene models as potential targets for pest control, we manually curated several gene families with a focus on genes that have key functional roles in *D. citri* biology and CLas interactions. This community effort produced 530 manually curated gene models across developmental, physiological, RNAi regulatory, and immunity-related pathways. As previously shown in the pea aphid, RNAi machinery genes putatively involved in the microRNA pathway have been specifically duplicated. A comprehensive transcriptome enabled us to identify a number of gene families that are either missing or misassembled in the draft genome. In order to develop biocuration as a training experience, we included undergraduate and graduate students from multiple institutions, as well as experienced annotators from the insect genomics research community. The resulting gene set (OGS v1.0) combines both automatically predicted and manually curated gene models. All data are available on https://citrusgreening.org/.

## INTRODUCTION

The Asian citrus psyllid (ACP), *Diaphorina citri* Kuwayama (Hemiptera:Liviidae), is a phloem-feeding insect native to Southeastern and Southwestern Asia with a host range limited to plants in the citrus genus and related Rutaceae spp. (1). Accidental anthropogenic introductions of psyllid-infested citrus combined with the ability of psyllids to disperse rapidly has allowed *D. citri* to extend its distribution to most of southern and eastern Asia, the Arabian Peninsula, the Caribbean, and South, Central and North America(1–6). For years, ACP has been classified as a global pest that is capable of devastating citrus crops through transmission of the bacterial agent, *Candidatus* Liberibacter asiaticus, CLas, which is associated with Huanglongbing (HLB) or citrus greening disease. The psyllid alone has little economic importance and causes only minor plant damage while feeding (7,8).

HLB is the most destructive and economically important disease of citrus, with practically all commercial citrus species and cultivars susceptible to CLas infection(9). Infected trees yield premature, bitter and misshapen fruit that is unmarketable. In addition, tree death follows 5-10 years after initial infection (2,9,10). Furthermore, HLB drastically suppresses economic progress in southern and eastern Asia by impeding viable commercial citrus agriculture within those regions (11). Florida is one of the top citrus-producing regions in the world and the largest in the United States, with nearly double the output of California, the second largest citrus-producing state (12). HLB puts the 9 billion dollar Florida citrus industry, with an annual net value of 1.5 billion dollars, at tremendous risk [USDA 2009]. In 2008, the HLB infection rate within central Florida was low (1.4% to 3.6%), but reaching 100% in the southern and eastern portions of the state (13,14). In 2005, when HLB was first detected in Florida, 9.3 million tons of oranges were harvested, but production has declined steadily to 5.3 million tons in 2016 as ACP and HLB have spread (12).

Primary management strategies focus on disrupting the HLB transmission pathway by suppressing psyllid populations and impeding interactions between CLas and psyllids. These strategies currently rely on extensive chemical application, which has broad environmental impact and high costs, and are ultimately unsustainable. To develop molecular methods that exploit current gene-targeting technologies, detailed genetic and genomic knowledge, including a high quality official gene set (OGS), is required (15,16). Early efforts focused on *D. citri* transcript expression(3, 17–19), analysis of the full transcriptome (20,21) and, more recently, analysis of the *D. citri* proteome (15). Arp et al. (22) performed a BLAST-based inventory of NCBI-predicted immune genes in *D. citri* (v100, see Methods). In contrast, we have conducted broad structural and functional annotation with the aid of a comprehensive transcriptome and created an official gene set for *D. citri* with a focus upon completing the repertoire of immune genes.

Manual curation improves the quality of gene annotation and establishing a ‘version controlled’ official gene set (OGS) provides a set of high quality, well-documented genes for the entire research community. Although ACP is a significant agricultural pest, it is not a model organism and the size of the research community does not warrant “museum” or "jamboree” annotation strategies (23). To maximize the number of genes annotated in a relatively short time, we augmented the “cottage industry” strategy (24) by training undergraduates to perform basic annotation tasks. The dispersed ACP annotation community agreed on a set of standard operating procedures and defined a set of primary gene targets. Starting with automated gene predictions, we used several additional types of evidence including RNAseq and proteomics data including comparisons to other insects to generate the first official gene set (OGS v1.0). Using an independent transcriptome enabled us to identify a number of gene families that are either missing or misassembled in the draft genome.

Our manual curation efforts focused on genes of potential use in vector control including immunity-related genes and pathways, RNAi machinery genes, multiple clans of cytochrome P450 genes and other genes relevant to insect development and physiology. We speculate that targeted analysis of these genes in *D. citri* will provide the foundation for a better understanding of the interactions between psyllid host and CLas pathogen, and will open the possibilities for research that can eventually find solutions to manage the dispersion of this very destructive pest and HLB.

## RESULTS AND DISCUSSION

### NCBI-Diaci 1.1 draft genome assembly

The Diaci1.1 draft genome assembly was generated using Illumina paired-end and mate-pair data with low coverage Pacbio for scaffolding and uploaded to NCBI (PRJNA251515) and Ag Data Commons (25) after filtering out bacterial contamination. Illumina sequencing was performed on the HiSeq2000 using 100bp or longer reads. Seven libraries were sequenced, with inserts ranging from “short” (ca. 275bp) to 10Kb. These are available in NCBI SRA and include 99.7 million paired-end reads (NCBI SRA: SRX057205), 35.1 million 2kb mate-pair reads (NCBI SRA: SRX057204), 30 million 5kb mate-pair reads (NCBI SRA: SRX058250) and 30 million 10kb mate-pair reads (NCBI SRA: SRX216330). A second round of DNA sequencing was performed with PacBio at 12X coverage (NCBI SRA: SRX218985) for scaffolding the Diaci1.0 Illumina assembly to create the Diaci 1.1 version of the *D. citri* genome. The Illumina data was assembled with the velvet (26) assembler followed by scaffolding with Pacbio long reads using the Pbjelly (27) pipeline (see Methods). The *D. citri* genome has an estimated size of 400-450Mb (18)(28). This genome has a length of 485 Mb with 19.3Mp of N’s. It contains 161,988 scaffolds with an N50 of 109.8kb. Given the high degree of fragmentation, we performed a Benchmarking sets of Universal Single-Copy Orthologs (BUSCO) (29) analysis with a set of conserved single-copy markers. A BUSCO analysis identifies the proportion of known single copy genes correctly assembled in a given dataset. The accuracy and resolution of this analysis is enhanced by using a set of markers specific to a phylogenetic clade so we used 3,550 markers from the nine insects in the Hemipteran order based on orthologous groups defined in the ORTHODB v9 (30) database. We found a significant number of these genes to be missing (35.7%, See Supplementary Table 1a) as confirmed in the curation section below.

**Table 1:** Immune gene pathways and gene counts in ACP, Pea and Aphid Whitefly. ACP genes identified only in MCOT v1.0 are in ().

|  | Pathway/Genes | ACP | Pea aphid | Whitefly |
| --- | --- | --- | --- | --- |
| *Pathogen recognition molecules* |  |  |  |  |
|  | CTLs: C-Type Lectins | 10 | 5 | 5 |
|  | GALEs: Galactoside-Binding Lectins | 4 | 1 | 4 |
|  | FREPs: Fibrinogen-Related Proteins | 3 | 1 | 5 |
|  | PGRPs: Peptidoglycan Recognition Proteins | 1 | 0 | 1 |
|  | BGBPs: 1,3-beta-D Glucan Binding Proteins | 0 | 2 | 5 |
|  | TEPs: Thio-Ester Containing Proteins | 2 | 1 | 1 |
| *Signaling cascades associated with pathogenesis* |  |  |  |  |
| Toll pathway and receptors | Toll receptors | 5 | 7 | 5 |
|  | Spaetzle | 5 | 6 | 9 |
|  | Tube | 1 | 1 | 1 |
|  | Pelle | 1 | 1 | 1 |
|  | MyD88 | 1 | 1 | 1 |
|  | CACT | 1 | 1 | 0 |
|  | TRAF6 | 1 | 2 | 1 |
| IMD pathway members | CASPAR | 3 | 1 | 1 |
|  | FADD | 0 | 0 | 0 |
|  | IKKB/ird5 | 1 | 0 | 0 |
|  | IMD | 0 | 0 | 0 |
|  | TAK1 | 1 | 1 | 1 |
|  | TAB | 1 | 1 | 1 |
| JAKSTAT pathway members | DOME | 1 | 3 | 1 |
|  | HOP | 1 | 1 | 1 |
|  | STAT | 1 | 3 | 1 |
| *Response to pathogens and pathogen-associated stress* |  |  |  |  |
|  | AMPs: Anti-Microbial Peptides | 0 | 7 | 4 |
|  | LYSs: Lysozymes | 5 | 3 | 5 |
|  | SODs: Superoxide Dismutases | 4 | 4 | 5 |
|  | CLIPs: CLIP-Domain Serine Proteases | 14 | 3 | 6 |
|  | Autophagy | 15 (2) | 8 | 16 |
|  | PPOs: Prophenoloxidasases | 2 | 2 | 4 |
|  | IAPs: Inhibitors of Apoptosis | 4 | 7 | 4 |

### MCOT transcriptome

To generate a more comprehensive set of gene models, we created the *D. citri* MCOT v1.0 transcriptome assembly. The MCOT pipeline (31) merges the output of multiple gene prediction and transcriptome assembly tools by clustering similar transcripts and selecting the best predicted protein for each cluster (see Methods). By supplementing genome-based transcript models with *de* novo-assembled transcripts, we hoped to obtain a more complete collection of *D. citri* transcripts. <u>M</u>aker v1.1 (32,33) gene models were predicted on the Diaci1.1 genome using RNAseq data from adult, nymph and egg tissue (20). The RNAseq data sets were also used to generate a genome-based transcriptome assembly using <u>C</u>ufflinks (34). *De novo* transcriptome assemblies of the adult, nymph and egg RNAseq data were performed with Oases (35) and Trinity (36). These data sets are all available at ftp://ftp.citrusgreening.org/annotation/MCOT/. *D.citri* MCOT v1.0 contains 30,562 proteins and is also available at Ag Data Commons (37).

The completeness of *D. citri* MCOT v1.0 was assessed with the BUSCO version 2 (29) tool with Hemipteran specific markers as described above. *D. citri* MCOT v1.0 contains 3,114/3,350 (92.9%) complete BUSCO orthologs, 2,239 (66.8%) of which are single copy and 875 (26.1%) appear to be duplicated. Four additional BUSCO orthologs (0.1%) are fragmented, and only 232 (7%) were not found. BUSCO analysis was also performed on the other resources used for annotation including three stage-specific *de novo* assembled transcriptomes from egg, nymph and adult tissue (20), the NCBI-Diaci1.1 genome assembly itself and the Maker v1.1 as well as the NCBI v100 predicted gene models (Figure 1, Supplementary Tables 1a and 1b). The *D.citri* MCOT v1.0 proteins that could be mapped to the genome assembly were also included in this analysis. The Maker v1.1 annotation set contains 18,205 protein-coding gene models. NCBI *D. citri* Annotation Release 100 (NCBI v100, https://www.ncbi.nlm.nih.gov/qenome/annotationeuk/Diaphorina/citri/100/) contains a total of 19,311 protein-coding gene models along with non-coding RNAs (776) and pseudogenes (207). As shown in Figure 1 and Supplementary Table 1b, none of these data sets proved to be as complete as *D.citri* MCOT v1.0, which contains a higher percentage of complete BUSCO orthologs and fewer fragmented or missing markers. Proteins representing transcripts assembled by Trinity (36) and Oases (35) exclusively from RNAseq data improve the completeness of MCOT v1.0 in comparison to the NCBI v100 annotation based on the fragmented and incomplete Diaci 1.1 draft genome.

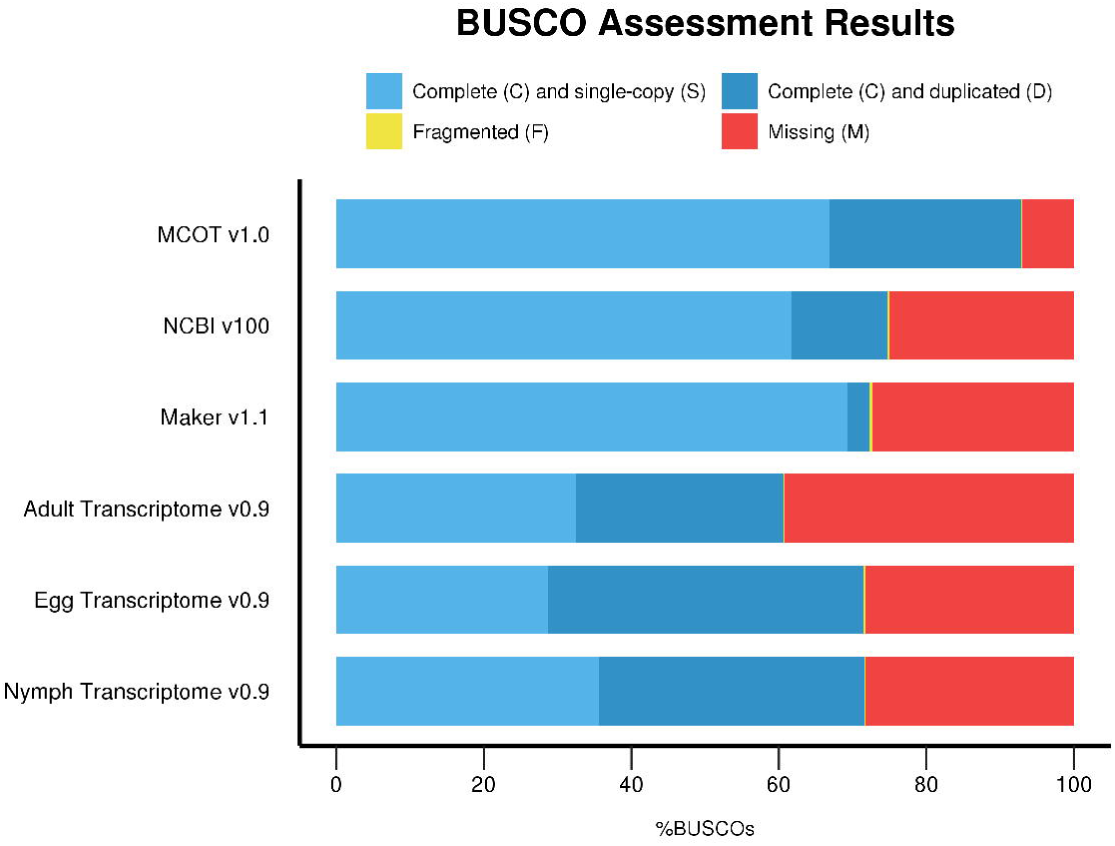

The functional annotation for MCOT v1.0 proteins was generated using InterproScan5, BLAST and AHRD (38), which assigned descriptions to 23,098 genes (75.6%), GO annotations to 15,314 genes (50%) and Pfam domains to 18,170 genes (59%). MCOT is available at ftp://ftp.citrusqreeninq.orq/annotation/MCOT/ and Ag Data Commons (37).

### Manual curation workflow

Manual curation of the *D. citri* (NCBI-Diaci1.1) draft genome assembly was undertaken to improve the quality of automated annotations (Supplementary Table 2) produced by the Maker (Maker v1.1) and NCBI pipelines (NCBI v100). This community-based manual annotation was focused on immunity-related genes as targets for ACP control.

**Table 2:** Homolog number of core machinery and auxiliary RNAi components in insects. Proteins highlighted in orange are RNase Type III enzymes. Proteins highlighted in blue are dsRNA binding proteins. Proteins highlighted in green are AGO family proteins. Proteins highlighted in yellow have been implicated in RISC or in small RNA biogenesis. *indicates homolog number was determined by BLAST and reciprocal BLAST analysis using NCBI’s non redundant databases. Homolog number with no asterisk were determined by publications or reported in ImmunoDB. ^†^ Indicates homolg is likely present but was unablr to be annotated in the current genome assembly. Note: Drosophila melanogaster has six other RNA helicase gene to Rm62. Some of the mosquito Rm62 proteins could be orthologous to these related RNA helicases.

|  | Dcr1 | Dcr2 | Drosha | Loqs | R2D2 | Pasha | AGO1 | AGO2 | AGO3 | PIWI/Aub | Armi | TSN | VIG-1 | Spn-E |
| --- | --- | --- | --- | --- | --- | --- | --- | --- | --- | --- | --- | --- | --- | --- |
| *D. melanogaster* | 1 | 1 | 1 | 1 | 1 | 1 | 1 | 1 | 1 | 2 | 1 | 1 | 1 | 1 |
| *A. gambiae* | 1 | 1 | 1 | 1 | 1 | 1 | 1 | 1 | 1 | 2 | 1 | 1 | 1 | 1 |
| *A. aegypti* | 1 | 1 | 1 | 1 | 1 | 1 | 2 | 1 | 1 | 7 | 2 | 1 | 1 | 1 |
| *C. quinquefasciatus* | 1 | 1 | 1 | 1 | 1 | 1 | 1 | 2 | 1 | 6 | 2 | 1 | 0* | 1 |
| *T. castaneum* | 1 | 1 | 1 | 1 | 2 | 1 | 1 | 2 | 1 | 1 | 3* | 1* | 2* | 1* |
| *C. lectularius* | 1 | 1 | 1 | 1 | 1 | 1 | 2 | 2 | 1 | 4 | 2* | 1* | 1* | 1* |
| *D. citri* | 1 | 1 | 1 or 2 | 2 | 1<sup>ψ</sup> | 2 | 1 | 1 | 1 | 1 | 1 | 2 | 1 | 2 |

| Species | Number of P450s |
| --- | --- |
| *Apolygus lucorum* (mirid bug) | 46 |
| *Acyrtosiphon pisum* (pea aphid) | 58 |
| *Cyrtorhinus lividipennis* (green miridbug) | 59 |
| *Cimex lectularius* (bedbug) | 60 |
| ***Diaphorina citri* (Asian citrus psyllid)** | 60 |
| *Laodelphax striatellus* (small brown planthopper) | 63 |
| *Nilaparvata lugens* (brown planthopper) | 66 |
| *Rhodnius prolixus* (kissing bug) | 87 |
| *Bemisia tabaci* (whitefly) | 128 |
| *Homalodisca vitripennis* (glassy winged sharpshooter) | 142 |

The Apollo Genome Annotation Editor hosted at i5k Workspace@NAL (https://i5k.nal.usda.gov/) was implemented for community-based curation of gene models (39). Multiple evidence tracks (Supplementary Table 3) were added to Apollo to assist in manual curation. Standard operating procedures were outlined at the beginning and refined based on feedback from annotators and availability of evidence resources. A typical workflow for manual gene curation involved selection of orthologs from related species, search of the ACP genome and MCOT v1.0 transcriptome, followed by assessment and correction, of the gene models based on evidence tracks to generate the final model. Any exceptions to this general workflow are specified in the respective gene reports (Supplementary Notes 1- 39). We used ImmunoDB (40) as a primary source of orthologs for curation of immune genes (Supplementary Notes 1-31). However, we also report other gene families of functional and evolutionary importance (Supplementary Notes 32-39) including aquaporins, cuticle proteins and secretory proteins. Following correction, final gene models were verified using reciprocal BLAST analysis. With this community-based curation effort, we annotated a total of 530 genes, the majority of which include genes predicted to function in immunity, development and physiology.

### Annotation Edit Distance (AED) as a measure of quality for different annotations

We employed the Annotation Edit Distance (AED) (33,41) metric to evaluate the quality of the different annotation data sets based on evidence from expression data. AED measures congruence of a predicted gene model with the RNAseq evidence supporting it. AED scores range between 0 and 1, where with an AED score of 0 denotes perfect concordance and an AED score of 1 denotes lack of any supporting evidence. The AED had a two-fold application here, selection of the best predicted gene set and quantification of improvement after manual curation. In order to generate AED scores, we compared the annotations to known insect proteins (NCBI taxonomy: “Hexapoda [6960]”) and to a comprehensive genome-guided transcriptome assembled from the latest data (See Methods and Supplementary Table 4) and the NCBI-Diaci1.1 draft assembly using the Maker pipeline (32). Most of the RNAseq data used to create this transcriptome was not used in the prediction of genes by other pipelines (NCBI v100, Maker v1.1 and MCOT v1.0) since it had not yet been produced and therefore provides independent validation. RNAseq data used for building the transcriptome included adult, nymph, egg, CLas exposed and healthy (adult and nymph tissue) as well as gut-specific expression data (see Supplementary Table 4). We calculated AED scores for NCBI v100, Maker v1.1 and mapped MCOT v1.0 annotation sets.

The plot for the AED cumulative fraction of transcripts (Figure 2) shows higher expression evidence support for NCBI v100 genes compared to the Maker v1.1 and mapped MCOT v1.0 annotation sets. Therefore the NCBI v100 annotations were selected to create the official gene set v1 (OGS v1.0). Although the MCOT v1.0 scores higher than NCBI v100 in BUSCO results (Figure 1 and Supplementary Table 1b), the mapped-MCOT set scores lower in the AED analysis as a large number of MCOT proteins could not be mapped on the draft Diaci 1.1 genome. The AED plot also shows that there are many gene models in all the annotation sets that do not have any expression evidence support (AED = 1.0). This could be due either to lack of RNAseq data for some of the loci or incorrect annotations. The predicted annotations may also have been affected by misassemblies in the NCBI-Diaci1.1 draft genome.

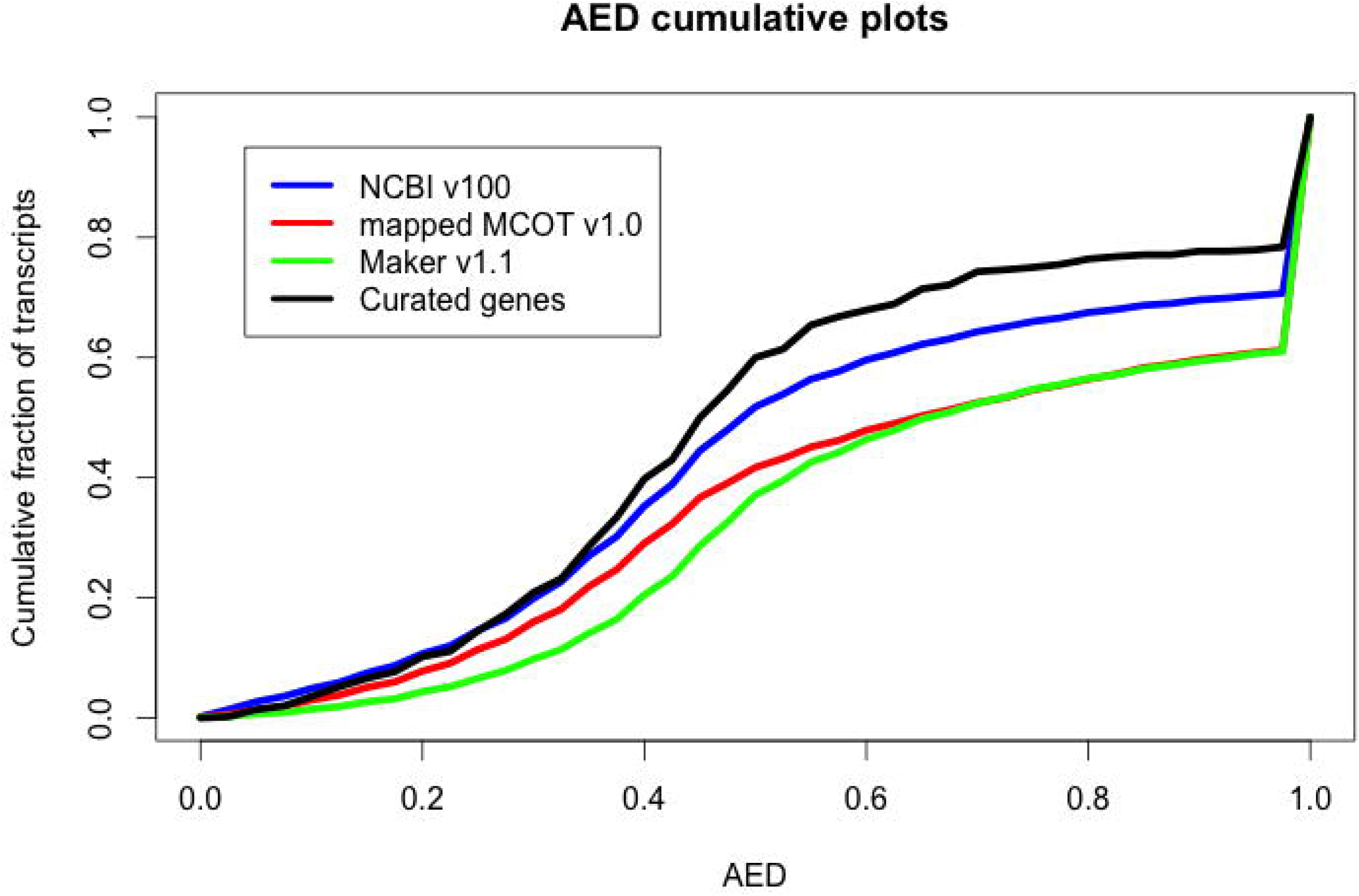

To quantify improvements in the gene structure by manual curation, we calculated AED scores for only the curated genes (530 genes). Cumulative fractions of transcripts of AED for curated genes show improvements indicating that the intron-exon structures in the curated genes have been corrected by manual curation (Figure 2).

### Training and curation strategy

#### Harnessing the Crowd

It is not possible for a single individual or computer system to fully curate a genome with precise biological fidelity. A growing number of genome sequencing projects have come to fruition thanks to the combined efforts of global consortia, (for example parasitoid wasps (42), centipede (43) and bed bug (44)). This indicates that a centralized model of genome annotation, the design used in earlier sequencing projects for the fruit fly (45) and the human genome (46), is giving way to a global and collaborative communities to generate high quality genome annotation. Beyond the problem of scale, curators require insights from others with expertise in specific gene families, which makes the process of curation inherently collaborative. Mobilizing groups of researchers to focus on these specific and manageable areas is more likely to distill the most pertinent and valuable knowledge from genome analysis.

We formed a team of 36 curators by enlisting collaborators primarily distributed across seven academic institutions (Indian River State College, Cornell University / Boyce Thompson Institute, Kansas State University, University of Cincinnati, University of Florida, University of Otago and University of Tennessee Health Center), including undergraduate and graduate students, postdocs, staff researchers, and faculty. The i5k Workspace@NAL (39) was selected as the platform for collaborative gene model curation as it is used widely by expert annotators from the insect genomics community. Training was provided at in-person workshop sessions and video conferences. The initial training workshop was structured to provide a review of principles associated with identification of specific genes by comparison to orthologs, gene structural aspects, how to search the genome for targets of interest, and using the combination of gene predictions and additional evidence tracks (Supplementary Table 3) to correct gene models with the Apollo platform. Lastly, we demonstrated secondary analyses, such as BLAST comparison to other insects and phylogenetic analyses, to the annotators to examine specific genes or gene sets. The training materials used for the workshop are included in supplementary data and also available online (47–49).

After initial in-person training, we continued to organize annotation via bi-weekly video conferences, documents on Google Drive and an online project management website (https://basecamp.com/2923904/projects/9184795). The project management website was used for coordinating all annotation activities and storage of working documents as well as all presentations and tutorials. The entire annotation workflow is shown in Figure 3 and a detailed training tutorial for the *D. citri* genome is included in supplementary data. The online forum facilitated discussions outside of the video conferences and allowed annotators to interact at their convenience. We established standard operating procedures for minimum evidence required for annotating a gene model, gene naming conventions and quality control checks (Supplementary Tables 5a and 5b). The default annotation workflow (Figure 3) was followed by majority of student annotators while the expert workflow was used by more experienced annotators.

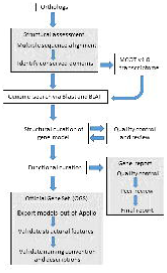

The undergraduate annotators at each site were mentored by a local faculty who coordinated separate in-person meetings. A few experienced annotators from the i5k community also participated in this curation effort. The video conferences facilitated a healthy discussion about curation methods among different groups of annotators. The combination of video conferences and online forum provided curators with the tools required to efficiently share data and information as they worked in teams across institutional boundaries. Moreover, since expert community curators volunteer their time to multiple projects, this model allowed them to contribute according to their availability.

#### Curation workflow

Gene lists and orthologs were made available via the project management website (https://basecamp.com/2923904/projects/9184795) so that the annotators could volunteer to curate genes of interest. We used ImmunoDB (40) as a primary source of immune gene orthologs, which provided expert-curated immunity genes for *Aedes aegypti, Anopheles gambiae, Drosophila melanogaster* and *Culex quinquefasciatus.* Other closely related organisms used to source of annotated orthologs included bed bug (*Cimex lectularius*) (44), pea aphid (*Acyrthosiphon pisum*)(*50*) and milkweed bug (51). All communication was done via emails and the project management website, which was also used to store presentations, gene reports, meeting minutes, presentations, working documents and data files. Annotation updates are shared with the community through the citrusgreening.org website (https://citrusgreening.org/annotation/index).

Candidate gene models were identified on the *D. citri* genome by using orthologous proteins as query in Apollo blat and i5k BLAST. The NCBI conserved domains database (52) was used to identify the conserved domains in the orthologs and candidate genes. Multiple sequence alignments were generated using MUSCLE(53), tcoffee (54) and clustal (55) to compare the ACP gene model to the query gene set. The final model was refined in Apollo using homology, RNAseq and proteomics evidence tracks. MEGA7 (56) was used to construct phylogenetic trees. Please see individual gene reports in supplementary notes 1-39 for detailed methods. When available, published literature was used to putatively assign molecular functions, participation in biological processes, and cellular localization for an annotated gene, associate term identifiers from the Gene Ontology Database (57) and PubMed identifiers from NCBI. Curated genes were assigned names and descriptions based on the function and domain structures available in published literature or NCBI.

Another strategy for identification of candidate genes followed by more experienced curators involved preprocessing the query before searching the ACP genome on Apollo. A set of representative genes was BLASTN searched against a database of all contigs in the ACP genome. These contigs were BLASTX searched against all the database of named insect genes. Analysis of the match helped to identify missing exons and extend partial exons to achieve the best possible manually curated version of the ACP gene. Once gene models were reconstructed from the genomic DNA, they were located on the *D. citri* genome using the i5k BLAST server (39). The models were then manually edited based on evidence tracks to produce the final gene model.

We performed multiple cycles of internal review of curated gene models to identify errors and suggest improvements to annotators. This was valuable in ensuring that annotators followed consistent guidelines throughout annotation. Curation of a gene family by an annotator was followed by a presentation summarizing their results during the video conference and peer-review. Annotations were ranked a scale of A-D (Supplementary Table 5a) during review depending on completeness and support from various evidence tracks. Standard gene naming conventions were defined and agreed upon to ensure consistency (Supplementary Table 5b).

The curated gene models were exported from Apollo and all functional annotations were manually checked for consistency with community standards (Supplementary Table 5b). The gene set was validated using the i5k quality control pipeline (https://github.com/NAL-i5K/I5KNALOGS/wiki/QC-phase) which identifies intra-model, inter-model and single-feature errors. The cleaned manual annotations were then merged with the protein-coding genes from the NCBI Diaphorina citri Annotation Release 100 (NCBI v100, ftp://ftp.ncbi.nlm.nih.gov/genomes/Diaphorina_citri/ARCHIVE/ANNOTATION_RELEASE.100/) using the NAL’s Merge prototype software (described at https://github.com/NAL-i5K/I5KNAL_OGS/wiki/Merge-phase; software is available on request). Non-coding RNAs from the NCBI Diaphorina citri Annotation Release 100 were added to the gene set after this merge, resulting in the Official Gene Set Dcitr_OGSv1.0 (58).

#### Education

Manual curation was incorporated into a bioinformatics class in 2016 by one of the authors (Benoit) at the University of Cincinnati. Specifically, one week in the class was utilized for genome annotation through the through the i5k genome annotation workspace for twenty-two students. This focused on how RNAseq datasets contribute to gene prediction and why these models need to be corrected manually before genome publication. Each student was responsible for correcting two gene models for *D. citri* in Apollo. As part of this course, students were given an assessment test before and after the class. Importantly, the rubric for this survey was not specifically designed for this project, rather the ACP genome was selected as the focus organism to annotate after the initial assessment test. Three questions (Supplementary Table 6, Details necessary for correct answers are included within this table) of this test focused on gene prediction and genome annotation, which were the major educational focus of the genome annotation week. The average score on these questions before the class was only ~38%, which improved to ~90% at the completion of this course. These scores indicated that the students had a much improved knowledge of genome assembly and annotation following this class.

Two of the participating institutions (Benoit, University of Cincinnati; D’Elia, Indian River State College) integrated the *D. citri* genome annotation into senior capstone courses. Students that participated in capstone projects at UC covered eight topic areas; aquaporins, acidic amino acid transporters, glycosphingolipid metabolism, glycolysis, histone binding, vitamin metabolism, vitamin transport, and Hox genes. With the addition of the capstone course, a total of 28 UC students participated in the manual curation process. Six students participated in capstone projects at IRSC during which they annotated 15 gene families. In total, 25 students directly participated in the manual curation process, contributing to 39 gene reports. This strategy reinforces the use of undergraduate students in gene annotation, as students show an increase in learning and produce scientific reports which contribute to peer-reviewed publications.

### Immune pathway in *D. citri*

Identification of the pathogen-induced immune components in ACP is critical for understanding and influencing the interaction between *D. citri* and CLas. The repertoire of immune genes is known to be a very diverse functional group and includes proteins that recognize infectious agents and initiate a signal, members of signal transduction pathways that relay the message to the nucleus, and the genes that are transcribed in response to infection. Below, we have briefly summarized our findings from manual structural and functional curation of immunity-related genes. Additional details are provided in gene reports for specific gene families and in the supplementary notes 1- 31.

### Pathogen recognition molecules

Recognition of infectious agents, the first step in immune defense, relies on a variety of cellular receptors including C-type lectins, Galectins, fibrinogen related proteins (FREP), Peptidoglycan recognition proteins (PGRP), Beta-1, 3-Glucan recognition proteins (*β*GRP) and thioester-containing proteins (TEP). These proteins recognize pathogen associated molecular patterns (PAMPs) which are associated with microbial cells (59). Most of these recognition molecules are widely conserved in insects, although the copy number often varies.

#### C-type Lectins

Ten C-type lectins (CTLs) carbohydrate binding receptors (60,61), were identified in *D. citri* (Supplementary Note 1). These include three oxidized low density lipoprotein receptor genes and genes encoding C-type Lectin 3, C-type Lectin 5, C-type Lectin 8, E selectin, Perlucin, Agglucetin subunit alpha and an selectin-like osteoblast derived protein. The number of CTLs in *D. citri* is comparable to the number found in bed bugs (11), pea aphids (6) and honeybees (10).

#### Galactoside-Binding Lectins

A total of three Galactoside-Binding Lectins (galectins) and one partial galectin were identified and manually curated within the ACP genome (Supplementary Note 2). Galectins bind β- galactose with their structurally similar carbohydrate-recognition domains (62), which can function alone or in clusters creating a β-sandwich structure without Ca^2+^ binding sites (63–65). Although we found more galectins in ACP than has been previously reported in the pea aphid (2 genes, (66)) and bed bug (1 gene, (44)), we did not observe a substantial lineage specific expansion as seen in Dipterans.

Fibrinogen related proteins

Similar to other hemipterans (pea aphid and bed bug), few fibrinogen related proteins (FREP) have been identified in ACP. Three complete FREPs were manually annotated (Scabrous, Angiopoietin and Tenascin) although partial un-annotatable FREP gene models were also detected. Like several of the other recognition molecule classes, the FREPs appear to have expanded in mosquitoes (67)(Supplementary Note 3). The suggestion that this expansion is related to blood feeding is consistent with the apparent absence in ACP of ficolin, tachylectins and aslectin, which are likely involved in detecting blood-borne parasites (67).

PGRP and *β*GRP

We identified one PGRP gene in *D. citri.* Insects have two classes of PGRPs: large (L) and small (S) (68). PRGP-L proteins recognize Gram-negative bacteria and activate the Imd pathway. PRGP-S genes interact with *β*GRPs such as GNBP to recognize components of Gram-positive bacteria and then activate the Toll pathway. Based on sequence similarity to other insect proteins, the *D. ctiri* PGRP protein seems to belong to the S class. We did not find any GNBP genes in *D. citri* (Supplementary Note 4). This is somewhat surprising, since GNBPs have been found in several hemipterans including pea aphids (66), bed bugs (44) and brown planthoppers (69).

Thioester containing proteins

Only two Thioester containing proteins (TEP) were identified in ACP (Supplementary Note 5), which is comparable to the number found in *Acrythosiphon pisum* and *Nasonia vitripennis.*

TEPs are members of an ancient protein family that includes vertebrate C complement and alpha-2-macroglobulin proteins (70). Insect TEPs seem to play a similar role to their vertebrate homologs, binding to invaders such as parasites or microbes, marking them for degradation, and they are upregulated by the JAK/STAT pathway during innate immune response (71).

### Signaling cascades associated with pathogenesis

Once a potential infection has been detected, a cellular response is initiated by signaling cascade. Typically, Gram-positive bacteria and fungi cause activation of the Toll pathway, while the Imd pathway responds to Gram-negative bacteria (72,73). The JAK/STAT pathway plays a role in several immune functions, including antiviral defense (74).

#### Toll Pathway

We identified four Toll receptors in *D. citri* (Supplementary Note 6). Comparison of the Toll receptors found in various insects suggests that there were six ancestral Toll receptors: *Toll-1, Toll-6, Toll-2/7, Toll-8, Toll-9* and *Toll-10* (75,76). Phylogenetic analysis indicates that the *D. citri* genes are orthologs of *Toll-1, Toll-6, Toll-7* and *Toll-8,* but orthologs of *Toll-9* and *Toll-10* were not found. Pea aphids and bed bugs have Toll receptors from every class but *Toll-9* (44,66). We found orthologs of five of the six Spätzle (Spz) ligand classes, including Spz1, Spz3, Spz4, Spz5 and Spz6 (Supplementary Note 7). The lack of Spz2 is not surprising since it has only been reported in Diptera and Hymenoptera. The downstream Toll pathway components are represented by single genes in most insects (77). Consistent with this, we identified single copy orthologs of *tube, pelle, MyD88,* TRAF6, *cactus* and *dorsal* (Supplementary Notes 8-13). Taken together, our findings suggest that the Toll pathway is largely conserved in *D. citri,* as it is in other insects.

#### Imd Pathway

As has been observed for several other hemipterans (66, 78–81), many components of the Imd pathway appear to be missing in *D. citri* (Supplementary Note 14). We were unable to identify orthologs of Dredd, FADD, Imd, IKKG, and Relish in either the assembled genome or the MCOT transcriptome. We did, however, find orthologs of pathway components IKKB, TAK1 and TAB, as well as FAF1/Caspar, a negative regulator of the pathway. The apparent loss of Imd pathway genes in many hemipterans has led to speculation that association with Gram-negative endosymbionts may have favored the loss of these genes (50,81), although it should be noted that organisms such as Wolbachia are also found in many insects with intact Imd pathways (82). Several Gram negative bacteria have been identified as *D. citri* symbionts, including *Wolbachia, Candidatus* Carsonella, *Candidatus* Profftella armaturae, and an as yet unidentified enteric bacteria closely related to *Klebsiella variicola* and *Salmonella enterica* (83–87). Given this information, it is tempting to speculate that the loss of many Imd pathway genes may, in fact, be associated with the ability of *D. citri* to acquire and harbor at least some of these Gram negative symbionts and might also be important for its ability to act as a carrier of CLas.

#### JAK/STAT Pathway

The Janus kinase/signal transducer of activators of transcription (JAK/STAT) pathway is a signaling pathway that provides direct communication between the membrane and nucleus (88). We identified genes encoding the major components of the JAK/STAT pathway, namely the orthologs of *domeless, hopscotch* and *marelle/STAT92E* (Supplementary Note 15). The JAK/STAT pathway is involved in many developmental processes, in addition to its role in immunity, and has been found in all sequenced insects to date, including other hemipterans.

### Response to pathogens and pathogen-associated stress

In response to infection, insect cells employ microbicidal compounds such as antimicrobial peptides (AMP), lysozymes and reactive oxygen species (ROS) to destroy invading cells and also activate tissue repair, wound healing and haematopoiesis processes. We searched the ACP genome for antimicrobial compounds (AMPs and lysozymes), the melanization-inducing Clip-domain serine proteases (CLIP), the protective superoxide dismutases (SOD), and autophagy-related genes.

#### Antimicrobial peptides

Although more than 250 antimicrobial peptides (AMPs) have been identified in insects, we searched the ACP genome and the MCOT transcriptome for ten classes of known AMPs without success. The AMPs investigated included attacin, cecropin, defensin, diptericin, drosocin, drosomycin, gambicin, holotricin, metchnikowin and thaumatin. Some of these AMPs appear to be widely conserved while others have only been identified in a limited number of species. Although defensins are one of the most widely conserved, ancient groups of AMPs (89), the absence of defensin in ACP is not unprecedented as its absence has also been reported the hemipteran *A. pisum* (66). While the pea aphid is lacking defensin (as well as most other previously identified insect AMPs) it does contain six thaumatin (antifungal) homologs. Despite its presence in the closely related pea aphid, we were unable to identify thaumatin in the ACP genome. It must be noted that absence of previously identified AMPs does not necessarily suggest absence of all AMPs. AMPs are an extremely large, diverse group of molecules often defined by structure and function rather than conserved motifs, making identification through comparative sequence analysis difficult. Additionally, most AMPs are probably yet to be identified and will need to be discovered through experimental work as opposed to orthologous searches based upon currently available sequence information.

#### Lysozymes

Five genes encoding lysozymes were found in the *D. citri* genome (Supplementary Note 16). Lysozymes hydrolyze bacterial peptidoglycan, disrupting cell walls and causing cell lysis. Many insects produce lysozymes, particularly c-type lysozymes, and secrete them into the hemolymph following bacterial infection. C-type lysozymes that commonly defend against Gram-positive bacteria have been reported in many different insect orders including Diptera, Hemiptera, and Lepidoptera (90). Although c-type lysozymes were not found in the initial search of the ACP genome, two c-type lysozyme transcripts were found in *D. citri* MCOT v1.0 and were subsequently used to identify these genes in the ACP genome. Additionally, three i-type lysozymes were identified in the ACP assembly.

Superoxide dismutases

Insect hemocytes can produce a burst of reactive oxygen species (ROS) to kill pathogens (91). Since ROS are also damaging to host cells, superoxide dismutases are necessary to detoxify ROS. We found a total of four superoxide dismutase (SOD) genes in *D. citri* (Supplementary Note 17). Similar to other insects (92–94), *D. citri* contains both CuZn and Mn SODs. One of the *D. citri* genes is an Mn SOD and the other three are CuZn SODs.

#### CLIP

Eleven Clip-domain serine proteases (CLIP), from four distinct evolutionary clades (CLIPA, CLIPB, CLIPC and CLIPD), were manually annotated in the *D. citri* genome and corresponding models identified in MCOT v1.0 (Supplementary Note 18). These clades are present as multigene families in insect genomes and function in the hemolymph in innate immune responses (95). In *Drosophila,* CLIPs are involved in melanization and the activation of the Toll pathway (96).

Autophagy

Using the *D. citri* genome and the MCOT gene set, we identified *D. citri* orthologs of autophagy-related genes known in *Drosophila* (Supplementary Note 19). Autophagy is the regulated breakdown of unnecessary or dysfunctional components of the cell. This process is highly conserved among all animals and is critical to the regulation of cell degradation and recycling of cellular components. The main pathway is macroautophagy, where specific cytoplasmic components are isolated from the remaining cell in a double-membraned vesicle called the autophagosome (97–99). We identified 17 out of 20 autophagy-related genes (Supplementary Note 19). There is only a single autophagy-related 8 gene in ACP gene sets compared to two for the *Drosophila* gene set, but this is common for non-Dipteran insects (98). Thus, as expected, psyllids have the required repertoire of autophagy-related genes to undergo macroautophagy.

In summary, there is a reduction in the number of immune recognition, signaling and response genes in *D. citri* compared to insects from Diptera. The reduction in the immunity genes is also observed in other hemipteran insect genomes such as *A. pisum* (66), *P. humanus* (78), *Bactericera cockerelli* (79), *R. prolixus* (80) and *B. tabaci* (81). The reduction of immunity-related genes in these insects has been attributed to association with their endosymbionts, which sometimes complement the immunity of the insects (100,101). In addition, this reduction in immune genes may be associated with insects that feed on nutritionally poor and relatively sterile food sources, such as blood and fluid from the xylem/phloem (78,100). However, bloodfeeding mosquitoes actually show an increase in immune genes and this expansion has been attributed to the likelihood of encountering pathogens in their food source. Arp et al. (22) pointed out additional inconsistencies with the diet hypothesis, including the presence of a full immune system in an insect species that develops in a sterile environment.

### RNA interference pathway in *D. citri*

The RNA interference (RNAi) pathway is a highly conserved, complex method of endogenous gene regulation and viral control mediated through short interfering RNAs (siRNAs), microRNAs (miRNAs), and piwiRNAs (piRNAs). While all of these small RNA molecules function to modulate or silence gene expression, the method of gene silencing and the biogenesis differs (102). In *Drosophila,* it appears that genes in the RNAi machinery have subfunctionalized to have roles in specific small RNA silencing pathways (103–109). While the RNAi machinery genes have been shown to be conserved across major taxa, functional studies in insects have been limited to a handful of Diptera. Investigating the complement of RNAi genes in *D. citri* may provide insight into the role that RNAi has on the immune response of phloem-feeding insects and could aid in better use of RNAi as a tool for pest management (110–112).

#### Core machinery

Class II (Drosha type) and class III (Dicer type) RNase III enzymes play an essential role in the biogenesis of small RNA molecules with Drosha and Dicer1 functioning to produce miRNAs and Dicer2 functioning to produce siRNAs (113–115). Our analysis of the *D. citri* genome revealed four possible loci with identity to insect Dicer proteins (Supplementary Note 20). However, further analysis of the MCOT transcriptome suggests that *D. citri* contains only one gene orthologous to *Dicer1* (MCOT05108.0.CO) and one gene orthologous to *Dicer2* (MCOT13562.0.CO). The remaining two loci are likely the result of genome fragmentation and misassembles. BLAST analysis of Drosha also identified multiple loci with homology to other insect Drosha proteins. MCOT transcriptome analysis indicates at least one or possibly two *drosha* homologs are present in *D. citri* (Supplementary Note 21).

dsRNA binding proteins act in concert with RNase III-type enzymes to bind and process precursor dsRNA molecules into small effector molecules (105,109,116,117). In some cases, these dsRNA binding proteins also function to load small RNA molecules into the RISC (118–120). In *Drosophila,* Pasha partners with Drosha (117), Loquacious (Loqs) partners with Dicer1 (109,116), and R2D2 partners with Dicer2 (105). In the *D. citri* genome, we identified two *pasha* homologs (Supplementary Note 22) and two *loqs* homologs but were initially unable to identify a true r2d2 ortholog (Supplementary Note 23) in the genome. The apparent absence of r2d2 was consistent with previous reports (22,121). However, a search of the MCOT (MCOT18647.0.CO) transcriptome identified a gene with similarity to R2D2 orthologs from bed bug, *Tribolium castaneum* and mosquitoes. While r2d2 is likely to be present in the *D. citri* genome, it is not annotatable given the limitations of the current assembly. Alternatively, if *r2d2* is missing from in *D.citri,* it is possible that one of the Loqs proteins identified functions in the RNAi pathway (121), as Loqs has been shown to associate with Dicer2 in both *Drosophila* and *Aedes aegypti* (122,123).

Argonaute (AGO) proteins present small RNA guide molecules to their complementary targets through silencing complexes and provide the ‘Slicer’ catalytic activity that is required for mRNA cleavage in some RNA silencing pathways (124–127). In *Drosophila* AGO1 is involved in the miRNA pathway, AGO2 is involved in silencing by siRNAs (103,108) and AGO3, PIWI and Aubergine (Aub) function in the piRNA pathway (104,106,128). In the *D.citri* genome, we have identified 4 *AGO* genes, *AGO1, AGO2, AGO3* and one gene corresponding to the PIW/Aub class of proteins (Supplementary Note 24).

Auxiliary (RISC and other) factors

A subset of other genes known to be involved in the function or regulation of the RNA-induced silencing complex (RISC) in *Drosophila* and other organisms have been identified and annotated in the *D.citri* genome. The genes identified include two *Tudor Staphylococcal Nucleases* (TSN, Supplementary Note 25), one *vasa-intronic gene* (*vig-1,* Supplementary Note 25), one *armitage (Armi*) gene (Supplementary Note 27), and one *Fragile X Mental Retardation 1* (*FMR1,* Supplementary Note 28) gene. Additionally, several more genes known to be involved in the biogenesis or function of small RNA molecules have been identified. These include two *spindle-E* genes (Supplementary Note 29), one *Rm62* gene (Supplementary Note 30) and one *Ran* gene (Supplementary Note 31).

In summary, the *D.citri* genome has a full complement of RNAi machinery genes. Duplications are more frequent in genes that have previously been associated with the miRNA pathway (*drosha, pasha* and *loqs*) as opposed to the RNAi or piRNA pathways. This is an interesting finding as the same result was found upon analysis of the pea aphid genome (129) but was not seen in the whitefly genome (81).

### Building the foundation for P450/Halloween genes targeted to reduce insect pests

Cytochrome P450s (CYPs) in eukaryotes are heme containing membrane bound enzymes that activate molecular oxygen via a mechanism involving a thiolate ligand to the heme iron. Usually this requires an electron donor protein, the NADPH cytochrome P450 reductase, in the ER or ferredoxin and ferredoxin reductase in the mitochondria (130,131). Insects have four deep branching clades on phylogenetic trees and this represents some losses during evolution as up to 11 clades are found in other animals. These are termed CYP2, CYP3, CYP4 and mitochondrial clans in P450 nomenclature (132). Most species have a tendency to expand P450s in one or more clans via tandem duplications. One interpretation of these P450 "blooms” is diversification to handle many related compounds from the environment that may be toxic or potential carbon sources.

*D. citri* in its current assembly has 60 P450 genes that are identified and named as distinct P450s. There are also numerous fragments named as partials. *Diaphorina* has a P450 bloom in the clusters CYP3172, CYP3174, CYP3175, CYP3176, CYP3178 in the CYP4 clan. There is another in the CYP3167 family in the CYP2 clan and a third that includes CYP6KA, CYP6KC and CYP6KD in the CYP3 clan. *D. citri* has 3 CYP4G genes.

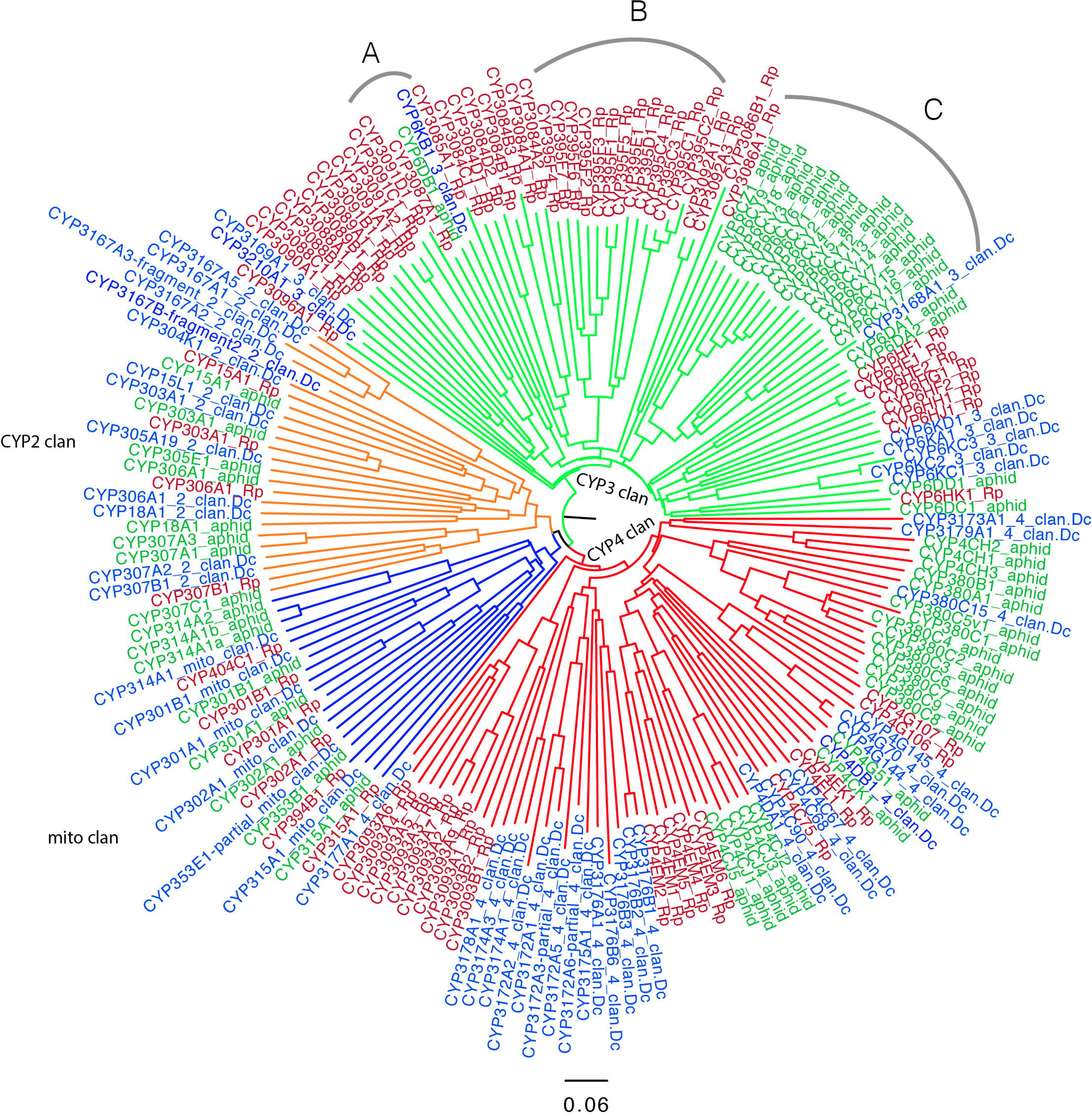

CYP2 and mito clans have many 1:1 orthologs but these are rare in the CYP3 and CYP4 clan (Figure 4). One exception in the CYP3 clan is CYP3087A1 and two neighbors CYP6DB1 and CYP6KB1 as they may be orthologs and probably should be in the same family. *R. prolixus* and *A. pisum* CYP3 clan genes have undergone gene blooms that were not found in *D. citri.* This may be interpreted as the common ancestor having few CYP3 clan P450s. The cluster at arc A (Figure 4) on the tree consisting of CYP6KB1, CYP6DB1 and CYP3087A1 may be evidence that these three are orthologs and should be in the same family. The large aphid clade of CYP6CY at arc C has no members from *Rhodnius* or *Diaphorina,* so it seems to be aphid specific. At arc B (Figure 4) the CYP395 family has four subfamilies C, D, E, F. There are many CYP395 genes in other hemipteran species, including *C. lectularius* (bedbug CYP395A,B), *Apolygus lucorum* (Hemiptera, a Mirid bug, CYP395G, H, J, K, L, M)), and *Cyrtorhinus lividipennis* (Hemiptera, green mirid bug, CYP395H, J, N). The fact they have been placed in different subfamilies suggests they are diverging from their common ancestor. The CYP3084 family with subfamilies A, B, C, D is only found in *Rhodnius* so far (133). The families CYP3088, CYP3089, CYP3090 and CYP3091 are also Hemiptera specific with some members in the same species noted earlier. The number of P450s varies with arthropod species from a low of 25 in the mite *Aculops lyoperscii* and 36 in *Pediculus humanus* (body louse) to over 200 in *Ixodes scapularis* (black-legged tick) and up to 158 in some mosquitos (134).

## CONCLUSION

We report the first draft assembly for the *D. citri* genome and the corresponding official gene set (OGS v1.0) which includes 530 manually curated genes and about 20,000 genes predicted by the NCBI Eukaryotic Genome Annotation Pipeline (NCBI v100). The community curation effort involved undergraduate students at multiple locations who were trained, individually or in a class setting, in gene curation as a part of this initiative. These students were supported by contributions from expert annotators in the insect genomics community. The major advantage of having both expert curators and undergraduate students work together in the annotation project was the training, exchange of ideas and community building. We also present standard operating procedures that can be used to guide and coordinate annotation by large virtual teams. We would like to note that implementing consistent annotation practices across a highly diverse and virtual team of annotators required regular discussions backed up by extensive documentation that was updated in response to user feedback. However, multiple rounds of manual review by senior annotators was still required to confirm that all annotations conform to certain basic criterion. A number of evidence sources (Supplementary Table 3) were added during the course of the project based upon availability and utility to ongoing annotation. One of the major decisions taken by the ACP community as a result of this annotation effort and subsequent detailed evaluation of the Diaci 1.1 genome was to generate an improved reference genome for ACP using the latest methods. An interim but improved assembly based on long read sequencing technology is available at Ag Data Commons (135).

This community annotation process will be continued by recruiting the next cohort of student annotators and scientists to improve structural and functional characterization of the next version of the ACP genome (135). The MCOT transcriptome reported in this paper is available at Ag Data Commons (37) and offers a genome-independent and comprehensive representation of the gene repertoire of *D. citri* that was used to improve the genome annotation. The MCOT transcriptome allowed us to identify lineage specific-gene models in *D. citri* and curate them. This gene set will support efforts in other hemipteran species that transmit bacterial pathogens.

In summary, we curated and described genes related to immunity and the RNAi pathway in addition to the cytochrome P450 genes. We report blooms in P450 genes in the CYP4, CYP2 and CYP3 clans which may be an evolutionary response to environmental stresses. Other important gene families that were curated as a part of the official gene set include aquaporins, cathepsins, cuticle and secretory proteins. We found the number of immunity-related genes to be reduced, even after direct targeting for improvement, in the *D. citri* genome similar to pea aphid and whitefly, which may reflect the association with microbial symbionts that have coevolved in both insects and the consumption of relatively sterile plant derived fluids. The genomic resources from this project will provide critical information underlying ACP biology that can be used to improve control of this pest.

MATERIAL AND METHODS

## DNA extraction and library preparation

High-molecular weight DNA was extracted using the BioRad AquaPure Genomic DNA isolation kit from fresh intact *D. citri* collected from a citrus grove in Ft. Pierce, FL and reared at the USDA, ARS, U.S. Horticultural Research Laboratory, Ft. Pierce, FL. To generate PacBio libraries, DNA was sheared using a Covaris g-Tube and SMRT-bell library was prepared using the 10Kb protocol (PacBio DNA template prep kit 2.0; 3-10Kb), cat #001-540-835.

## Genome sequencing and assembly

Samples were prepared for Illumina sequencing using the TruSeq DNA library preparation kits for paired-end as well as long-insert mate-pair libraries. Thirty-nine SMRTcells of the library were sequenced, all with 2×45 minute movies. A total of 2,750,690 post-filter reads were generated, with an average of 70,530 reads per SMRTcell. The post-filter mean read length was 2,504 bp with an error rate of 15%.

Velvet (26) was used with kmer 59 for generating the Diaci1.0 draft assembly. PacBio long reads were mapped to the draft assembly using blasr (136) with the following parameters: - minMatch 8 -minPctIdentity 70 -bestn 5 -nCandidates 30 -maxScore −500 -nproc 8 - noSplitSubreads. These alignments were parsed using PBJelly (27) with default parameters to scaffold the draft assembly and create the final Diaci 1.1 reference genome. More details about the history of ACP genome sequencing can be found at citrusgreening.org (https://citrusqreeninq.org/orqanism/Diaphorina_citri/genome). The Diaci 1.1 assembly was evaluated with BUSCO version 2 and the Hemipteran marker set with default parameters.

## Maker and NCBI annotation

The maker control files are included in supplementary data. BLAST 2.2.27+ was used with augustus version 2.5.5 (137) and exonerate version 2.2.0 (138). Only contigs longer than 10kb were selected for annotation with psyllid specific RNAseq data from Reese et al. (20). Details of ACP genome annotation by the NCBI Eukaryotic Genome Annotation pipeline are available at https://www.ncbi.nlm.nih.gov/genome/annotation_euk/Diaphorina_citri/100/.

## MCOT transcriptome assembly

Reads were assembled with Trinity in two runs, one used reads as single end reads, and the other used them as paired end reads. Reads were trimmed based on fastq quality score (the -- trimmomatic option was enabled and run under the default setting of Trinity). The transcripts from both runs were compiled together to create the final Trinity assembly.

Velvet-Oases (35) assemblies were performed for trimmed reads (trimmed in Trinity run, the read quality control step) from the egg, nymph and adult separately, with kmer length of 23, 25, 27 and 29 as single end reads, and kmer length 25 as paired end reads. The 15 outputs were combined using the Oases merge function (--long, kmer 27 and -min_trans_lgth 200) to generate the final assembly.

Reads from the egg, nymph and adult were first aligned to the genome with Tophat (139) with the insert length parameter based on each library (53, 24 and 90). The parameter --read-realign-edit-dist were set to 0 to ensure better alignment results. Gene models were generated by Cufflinks (140) with default settings, with the -frag-bias-correct and -multi-read-correct function (-b, - u) enabled to give the most accurate gene models.

Transcripts from Maker (32,33), Cufflinks (34), Oases (35)and Trinity (36) were translated to proteins with Transdecoder version 2.0.1(141) and only unique proteins were retained. Furthermore, protein sequences from each program (Maker, Cufflinks, Oases, Trinity) were compared using BLASTP with a special scoring matrix (matching score of non-identical amino acids set to -100 of the BLOSUM62 matrix) to proteins from other arthropod species. The best protein models from each source were selected to create the final MCOT v1.0 protein set, and the corresponding transcript set. MCOT v1.0 set has 30,562 genes and is available at ftp://ftp.citrusqreeninq.orq/annotation/MCOT/ and Ag Data Commons (37).

MCOT v1.0 proteins were analyzed using Interproscan 5 (142) based on InterPro databases with the options -goterms to get GO terms, -iprlookup to switch on look-up of corresponding InterPro annotation and -pa option to switch on lookup of corresponding pathway annotation.

We also performed BLASTP comparison (parameters: -e 0.0001 -v 200 -b 200 -m0) of the MCOT proteins to insect proteins from Uniprot and NCBI nr databases. These BLAST results were used as input for AHRD (Automated assignment of Human Readable Descriptions) (38), to assign functional descriptions to each MCOT protein. This functional annotation was performed using a filter of bit score of more than 50 and e-value less than e-10. Pfam domains and eene ontology terms were also assigned from the Interproscan analysis.

## Annotation edit distance

We mapped transcripts from MCOT v1.0 to the genome using GMAP (143). Out of 30,562 transcripts, 19,744 were mapped with at least 90% query coverage and 90% identity.

A genome guided transcriptome was generated to validate all annotation sets using all available RNAseq data (Supplementary Table 4) and insect proteins (NCBI taxonomy: “Hexapoda [6960]”) from Swiss-Prot were used as sources of evidence. 622 million paired-end reads were mapped to NCBI-Diaci1.1 assembly using hisat2 (144) with a mapping rate of 81.78%. Mapped files were sorted with samtools rocksort. After sorting, a genome-guided transcriptome assembly was performed using StringTie (145) and the resulting transcriptome contained 210,890 transcripts (N50: 1,691bp).

Annotation edit distance was calculated for all gene models from NCBI v100, mapped MCOT, Maker v1.1 and curated gene sets. The Maker genome annotation (v2.31.8) pipeline was used for calculating AED (33,41).

## ACKNOWLEDGEMENTS

Kascha Bohnenblust from Kansas State University assisted with the organization of meetings for the community curation effort. The following personnel at Los Alamos National Laboratory, Los Alamos, NM contributed to the Illumina and Pacbio sequencing: Kimberly McMurry, Cheryl D Gleasner, Krista Reitenga, Shunsheng (Cliff) Han and Goutam Gupta. Christian Haudenschild from lllumina, Inc. supported the genome assembly. The following personnel at USDA-ARS, U.S. Horticultural Research Laboratory, Fort Pierce, FL contributed to the sample preparation for genome sequencing: Maria T. Gonzalez, Belkis Diego and Kathy Moulton. We would like to thank Felipe Simāo for providing the Hemipteran marker set for BUSCO analysis.

## REFERENCES

1. Halbert, S. E. and Núñez, C. A. (2004) Florida Entomol., 87, 401–402, DISTRIBUTION OF THE ASIAN CITRUS PSYLLID, DIAPHORINA CITRI KUWAYAMA (RHYNCHOTA: PSYLLIDAE) IN THE CARIBBEAN BASIN.

2. Halbert, S. E. and Manjunath, K. L. (2004) Florida Entomol., 87, 330–353, ASIAN CITRUS PSYLLIDS (STERNORRHYNCHA: PSYLLIDAE) AND GREENING DISEASE OF CITRUS: A LITERATURE REVIEW AND ASSESSMENT OF RISK IN FLORIDA.

3. Boykin, L. M., De Barro, P., Hall, D. G., et al. (2012) Bull. Entomol. Res., 102, 573–582, Overview of worldwide diversity of Diaphorina citri Kuwayama mitochondrial cytochrome oxidase 1 haplotypes: two Old World lineages and a New World invasion.

4. French, J. V, Kahlke, C. J. and Da Graça, J. V (2001) Subtrop. Plant Sci., 53, 14–15, First record of the Asian citrus psylla, Diaphorina citri Kuwayama (Homoptera: Psyllidae) in Texas.

5. Pluke, R. W. H., Qureshi, J. A. and Stansly, P. A. (2008) Florida Entomol., 91, 36–42, CITRUS FLUSHING PATTERNS, DIAPHORINA CITRI (HEMIPTERA: PSYLLIDAE) POPULATIONS AND PARASITISM BY TAMARIXIA RADIATA (HYMENOPTERA: EULOPHIDAE) IN PUERTO RICO.

6. Tsai, J. H. and Liu, Y. H. (2000) J. Econ. Entomol., 93, Biology of Diaphorina citri (Homoptera: Psyllidae) on Four Host Plants.

7. Teixeira, D. do C., Saillard, C., Eveillard, S., et al. (2005) Int. J. Syst. Evol. Microbiol., 55, 1857–62, “Candidatus Liberibacter americanus”, associated with citrus huanglongbing (greening disease) in São Paulo State, Brazil.

8. Capoor, S. P., Rao, D. G., Viswanath, S. M., et al. (1967) Indian J. Agric. Sci., 37, 572–575, Diaphorina citri Kuway., a vector of the greening disease of citrus in India.

9. Bové, J. M. (2006) J. Plant Pathol 88, 7–37, I NVITED R EVIEW HUANGLONGBINGL: A DESTRUCTIVE, NEWLY-EMERGING, CENTURY-OLD DISEASE OF CITRUS 1.

10. Manjunath, K. L., Halbert, S. E., Ramadugu, C., et al. (2008) Phytopathology, 98, 387–396, Detection of’Candidatus Liberibacter asiaticus’ in Diaphorina citri and its importance in the management of citrus huanglongbing in Florida.

11. Leong, S. C. T., Abang, F., Beattie, A., et al. (2012) Sci. World J., 2012, Impacts of horticultural mineral oils and two insecticide practices on population fluctuation of Diaphorina citri and spread of huanglongbing in a citrus orchard in Sarawak.

12. Honig, L. October Crop Production Executive Summary https://www.nass.usda.gov/Newsroom/Executive_Briefings/2016/10_12_2016.pdf.

13. Tiwari, S., Lewis-Rosenblum, H., Pelz-Stelinski, K., et al. (2010) J. Econ. Entomol., 103, Incidence of Candidatus Liberibacter asiaticus Infection in Abandoned Citrus Occurring in Proximity to Commercially Managed Groves.

14. Tabachnick, W. J. (2015) J. Econ. Entomol., 108, 839–848, Diaphorina citri (Hemiptera: Liviidae) vector competence for the citrus greening pathogen Candidatus Liberibacter asiaticus.

15. Ramsey, J. S., Johnson, R. S., Hoki, J. S., et al. (2015) PLoS One, 10, e0140826, Metabolic Interplay between the Asian Citrus Psyllid and Its Profftella Symbiont: An Achilles’ Heel of the Citrus Greening Insect Vector.

16. de Andrade, E. C. and Hunter, W. B. In RNA Interference; InTech, 2016.

17. Marutani-Hert, M., Hunter, W. B. and Hall, D. G. (2010) Florida Entomol., 93, 519–525, Gene response to stress in the Asian citrus psyllid (Hemiptera: Psyllidae).

18. Hunter, W. B., Dowd, S. E., Katsar, C. S., et al. (2009) Gene, 18–29, Psyllid Biology: Expressed Genes in Adult Asian Citrus Psyllids, Diaphorina citri Kuwayama.

19. Hunter, W. B., Hail, D., Tipping, C., et al. In Symposium Proceedings 2010.

20. Reese, J., Christenson, M. K., Leng, N., et al. (2014) J Genomics, 2, 54–58, Characterization of the Asian Citrus Psyllid Transcriptome.

21. Fisher, T., Vyas, M., He, R., et al. (2014) Pathogens, 3, 875–907, Comparison of Potato and Asian Citrus Psyllid Adult and Nymph Transcriptomes Identified Vector Transcripts with Potential Involvement in Circulative, Propagative Liberibacter Transmission.

22. Arp, A. P., Pelz-Stelinski, K. and Hunter, W. (2016) Front. Physiol., 7, 570, Annotation of the Asian citrus psyllid genome reveals a reduced innate immune system.

23. Stein, L. (2001) Nat. Rev. Genet., 2, 493–503, Genome annotation: from sequence to biology.

24. Elsik, C. G., Worley, K. C., Zhang, L., et al. (2006) Genome Res., 16, 1329–33, Community annotation: procedures, protocols, and supporting tools.

25. Leng, N., English, A., Johnson, S., et al. Diaphorina citri genome assembly Diaci 1.1 2017.

26. Zerbino, D. R. and Birney, E. (2008) Genome Res., 18, 821–9, Velvet: algorithms for de novo short read assembly using de Bruijn graphs.

27. English, A. C., Richards, S., Han, Y., et al. (2012) PLoS One, 7, e47768, Mind the Gap: Upgrading Genomes with Pacific Biosciences RS Long-Read Sequencing Technology.

28. Marutani-Hert, M., Hunter, W. B. and Hall, D. G. In Vitro Cell. Dev. Biol. Anim., 45, 317–20, Establishment of Asian citrus psyllid (Diaphorina citri) primary cultures.

29. Simao, F. A., Waterhouse, R. M., Ioannidis, P., et al. (2015) Bioinformatics, btv351-, BUSCO: assessing genome assembly and annotation completeness with single-copy orthologs.

30. Waterhouse, R. M., Zdobnov, E. M., Tegenfeldt, F., et al. (2011) Nucleic Acids Res., 39, D283–8, OrthoDB: the hierarchical catalog of eukaryotic orthologs in 2011.

31. Cao, X. and Jiang, H. (2015) Insect Biochem. Mol. Biol., Integrated modeling of proteincoding genes in the Manduca sexta genome using RNA-Seq data from the biochemical model insect.

32. Cantarel, B. L., Korf, I., Robb, S. M. C., et al. (2008) Genome Res., 18, 188–96, MAKER: an easy-to-use annotation pipeline designed for emerging model organism genomes.

33. Yandell, M. and Ence, D. (2012) Nat. Rev. Genet., 13, 329–342, A beginner’s guide to eukaryotic genome annotation.

34. Trapnell, C., Williams, B. A., Pertea, G., et al. (2010) Nat. Biotechnol., 28, 511–5, Transcript assembly and quantification by RNA-Seq reveals unannotated transcripts and isoform switching during cell differentiation.

35. Schulz, M. H., Zerbino, D. R., Vingron, M., et al. (2012) Bioinformatics, 28, 1086–1092, Oases: robust de novo RNA-seq assembly across the dynamic range of expression levels.

36. Grabherr, M. G., Haas, B. J., Yassour, M., et al. (2011) Nat. Biotechnol., 29, Full-length transcriptome assembly from RNA-Seq data without a reference genome.

37. Saha, S., Cao, X., Flores, M., et al. Diaphorina citri MCOT transcriptome 2017.

38. Schoof, H. In Plant and Animal Genome XXIV Conference.

39. Poelchau, M., Childers, C., Moore, G., et al. (2014) Nucleic Acids Res., gku983-, The i5k Workspace@NAL--enabling genomic data access, visualization and curation of arthropod genomes.

40. Waterhouse, R. M., Kriventseva, E. V, Meister, S., et al. (2007) Science, 316, 1738–43, Evolutionary dynamics of immune-related genes and pathways in disease-vector mosquitoes.

41. Eilbeck, K., Moore, B., Holt, C., et al. Quantitative measures for the management and comparison of annotated genomes. 2009; Vol. 10.

42. Werren, J. H., Richards, S., Desjardins, C. A., et al. (2010) Science, 327, 343–8, Functional and evolutionary insights from the genomes of three parasitoid Nasonia species.

43. Chipman, A. D., Ferrier, D. E. K., Brena, C., et al. (2014) PLoS Biol., 12, e1002005, The First Myriapod Genome Sequence Reveals Conservative Arthropod Gene Content and Genome Organisation in the Centipede Strigamia maritima.

44. Benoit, J. B., Adelman, Z. N., Reinhardt, K., et al. (2016) Nat. Commun., 7, 10165, Unique features of a global human ectoparasite identified through sequencing of the bed bug genome.

45. Adams, M. D., Celniker, S. E., Holt, R. A., et al. (2000) Science (80-.)., 287, 2185–2195, The genome sequence of Drosophila melanogaster.

46. Venter, J. C., Adams, M. D., Myers, E. W., et al. (2001) Science (80-.)., 291, 1304–1351, The Sequence of the Human Genome.

47. Munoz-Torres, M. Apollo Workshop at KSU 2015 https://www.slideshare.net/MonicaMunozTorres/apollo-workshop-at-ksu-2015 (accessed Jan 1, 2017).

48. Munoz-Torres, M. Apollo Exercises Kansas State University 2015 https://www.slideshare.net/MonicaMunozTorres/apollo-exercises-kansas-state-university-2015 (accessed Jan 1,2017).

49. Munoz-Torres, M. Apollo annotation guidelines for i5k projects Diaphorina citri https://www.slideshare.net/MonicaMunozTorres/apollo-annotation-guidelines-for-i5k-projects-diaphorina-citri (accessed Jan 1, 2017).

50. Consortium, T. I. A. G. (2010) PLoS Biol., 8, e1000313, Genome sequence of the pea aphid Acyrthosiphon pisum.

51. Vargas Jentzsch, I. M., Hughes, D. S. T. and Poelchau, M. F. T. The O. fasciatus curation community, Richards S, Panfilio KA. 2015. Oncopeltus fasciatus official gene set v1. 1.

52. Marchler-Bauer, A., Derbyshire, M. K., Gonzales, N. R., et al. (2015) Nucleic Acids Res., 43, D222–6, CDD: NCBI’s conserved domain database.

53. Edgar, R. C. (2004) Nucleic Acids Res., 32, 1792–1797, MUSCLE: multiple sequence alignment with high accuracy and high throughput.

54. Notredame, C., Higgins, D. G. and Heringa, J. (2000) J. Mol. Biol., 302, 205–217, T-Coffee: A novel method for fast and accurate multiple sequence alignment.

55. Sievers, F., Wilm, A., Dineen, D., et al. (2011) Mol. Syst. Biol., 7, 539, Fast, scalable generation of high-quality protein multiple sequence alignments using Clustal Omega.

56. Kumar, S., Stecher, G. and Tamura, K. (2016) Mol. Biol. Evol., msw054, MEGA7: Molecular Evolutionary Genetics Analysis version 7.0 for bigger datasets.

57. Consortium, G. O. and others (2004) Nucleic Acids Res., 32, D258--D261, The Gene Ontology (GO) database and informatics resource.

58. Saha, S. Diaphorina citri Official Gene Set v1.0 2017.

59. Christophides, G. K., Vlachou, D. and Kafatos, F. C. (2004) Immunol Rev, 198, 127–148, Comparative and functional genomics of the innate immune system in the malaria vector Anopheles gambiae.

60. Dodd, R. B. and Drickamer, K. (2001) Glycobiology, 11, 71R--79R, Lectin-like proteins in model organisms: implications for evolution of carbohydrate-binding activity.

61. Cambi, A. and Figdor, C. G. (2003) Curr. Opin. Cell Biol., 15, 539–546, Dual function of C-type lectin-like receptors in the immune system.

62. Cummings, R. D. and Liu, F. T. Galectins Essentials of Glycobiology. 2. Vol. Chapter 33 2009.

63. Leffler, H., Carlsson, S., Hedlund, M., et al. (2002) Glycoconj. J., 19, 433–440, Introduction to galectins.

64. Mitchell, D. A., Fadden, A. J. and Drickamer, K. (2001) J. Biol. Chem., 276, 28939–28945, A novel mechanism of carbohydrate recognition by the C-type lectins DC-SIGN and DC-SIGNR Subunit organization and binding to multivalent ligands.

65. Wang, L., Wang, L., Yang, J., et al. (2012) Dev. Comp. Immunol., 36, 591–601, A multi-CRD C-type lectin with broad recognition spectrum and cellular adhesion from Argopectenirradians.

66. Gerardo, N. M., Altincicek, B., Anselme, C., et al. (2010) Genome Biol., 11, R21, Immunity and other defenses in pea aphids, Acyrthosiphon pisum.

67. Wang, X., Zhao, Q. and Christensen, B. M. (2005) BMC Genomics, 6, 1, Identification and characterization of the fibrinogen-like domain of fibrinogen-related proteins in the mosquito, Anopheles gambiae, and the fruitfly, Drosophila melanogaster, genomes.

68. Dziarski, R. and Gupta, D. (2006) Genome Biol., 7, 1, The peptidoglycan recognition proteins (PGRPs).

69. Bao, Y.-Y., Qu, L.-Y., Zhao, D., et al. (2013) BMC Genomics, 14, 1, The genome-and transcriptome-wide analysis of innate immunity in the brown planthopper, Nilaparvata lugens.

70. Blandin, S. and Levashina, E. A. (2004) Mol. Immunol., 40, 903–908, Thioester-containing proteins and insect immunity.

71. Agaisse, H. and Perrimon, N. (2004) Immunol. Rev., 198, 72–82, The roles of JAK/STAT signaling in Drosophila immune responses.

72. Lindsay, S. A. and Wasserman, S. A. (2014) Dev. Comp. Immunol., 42, 16–24, Conventional and non-conventional Drosophila Toll signaling.

73. Myllymäki, H., Valanne, S. and Ramet, M. (2014) J. Immunol., 192, 3455–3462, The Drosophila imd signaling pathway.

74. Myllymäki, H. and Rämet, M. (2014) Scand. J. Immunol., 79, 377–385, JAK/STAT pathway in Drosophila immunity.

75. Evans, J. D., Aronstein, K., Chen, Y. P., et al. (2006) Insect Mol. Biol., 15, 645–656, Immune pathways and defence mechanisms in honey bees Apis mellifera.

76. Benton, M. A., Pechmann, M., Frey, N., et al. (2016) Curr. Biol., Toll Genes Have an Ancestral Role in Axis Elongation.

77. Viljakainen, L. (2015) Brief. Funct. Genomics, elv002-, Evolutionary genetics of insect innate immunity.

78. Kim, J. H., Min, J. S., Kang, J. S., et al. (2011) Insect Biochem. Mol. Biol., 41, 332–339, Comparison of the humoral and cellular immune responses between body and head lice following bacterial challenge.

79. Nachappa, P., Levy, J. and Tamborindeguy, C. (2012) Mol. Genet. genomics, 287, 803–817, Transcriptome analyses of Bactericera cockerelli adults in response to Candidatus Liberibacter solanacearum infection.

80. Mesquita, R. D., Vionette-Amaral, R. J., Lowenberger, C., et al. (2015) Proc. Natl. Acad. Sci., 112, 14936–14941, Genome of Rhodnius prolixus, an insect vector of Chagas disease, reveals unique adaptations to hematophagy and parasite infection.

81. Chen, W., Hasegawa, D. K., Kaur, N., et al. (2016) BMC Biol., 14, 110, The draft genome of whitefly Bemisia tabaci MEAM1, a global crop pest, provides novel insights into virus transmission, host adaptation, and insecticide resistance.

82. Zug, R. and Hammerstein, P. (2012) PLoS One, 7, e38544, Still a Host of Hosts for Wolbachia: Analysis of Recent Data Suggests That 40% of Terrestrial Arthropod Species Are Infected.

83. Nakabachi, A., Yamashita, A., Toh, H., et al. (2006) Science (80-.)., 314, 267, The 160- kilobase genome of the bacterial endosymbiont Carsonella.

84. Hilgenboecker, K., Hammerstein, P., Schlattmann, P., et al. (2008) FEMS Microbiol Lett, 281, 215–220, How many species are infected with Wolbachia?--A statistical analysis of current data.

85. Saha, S., Hunter, W. B., Reese, J., et al. (2012) PLoS One, 7, e50067, Survey of Endosymbionts in the Diaphorina citri Metagenome and Assembly of a Wolbachia wDi Draft Genome.

86. Sloan, D. B. and Moran, N. A. (2012) Mol Biol Evol, 29, 3781–3792, Genome reduction and co-evolution between the primary and secondary bacterial symbionts of psyllids.

87. Nakabachi, A., Ueoka, R., Oshima, K., et al. Defensive Bacteriome Symbiont with a Drastically Reduced Genome; 2013; Vol. 23.

88. O’Shea, J. J., Schwartz, D. M., Villarino, A. V, et al. (2015) Annu. Rev. Med., 66, 311–328, The JAK-STAT pathway: impact on human disease and therapeutic intervention*.

89. Zhang, L. and Gallo, R. L. (2016) Curr. Biol., 26, R14--R19, Antimicrobial peptides.

90. Callewaert, L. and Michiels, C. W. (2010) J. Biosci., 35, 127–160, Lysozymes in the animal kingdom.

91. Lavine, M. D. and Strand, M. R. (2002) Insect Biochem. Mol. Biol., 32, 1295–1309, Insect hemocytes and their role in immunity.

92. Bordo, D., Djinovic, K. and Bolognesi, M. (1994) J. Mol. Biol., 238, 366–386, Conserved patterns in the Cu, Zn superoxide dismutase family.

93. Parker, J. D., Parker, K. M., Sohal, B. H., et al. (2004) Proc. Natl. Acad. Sci. U. S. A., 101, 3486–3489, Decreased expression of Cu--Zn superoxide dismutase 1 in ants with extreme lifespan.

94. Colinet, D., Cazes, D., Belghazi, M., et al. (2011) J. Biol. Chem., 286, 40110–40121, Extracellular superoxide dismutase in insects characterization, function, and interspecific variation in parasitoid wasp venom.

95. Kanost, M. R. and Jiang, H. (2015) Curr. Opin. insect Sci., 11,47–55, Clip-domain serine proteases as immune factors in insect hemolymph.

96. Veillard, F., Troxler, L. and Reichhart, J.-M. (2016) Biochimie, 122, 255–269, Drosophila melanogaster clip-domain serine proteases: Structure, function and regulation.

97. Chang, Y.-Y. and Neufeld, T. P. (2010) FEBS Lett., 584, 1342–1349, Autophagy takes flight in Drosophila.

98. Malagoli, D., Abdalla, F. C., Cao, Y., et al. (2010) Autophagy, 6, 575–588, Autophagy and its physiological relevance in arthropods: current knowledge and perspectives.

99. Zirin, J. and Perrimon, N. In Seminars in immunopathology; 2010; Vol. 32, pp. 363–372.

100. Altincicek, B., Gross, J. and Vilcinskas, A. (2008) Insect Mol. Biol., 17, 711–716, Wounding-mediated gene expression and accelerated viviparous reproduction of the pea aphid Acyrthosiphon pisum.

101. Ratzka, C., Gross, R. and Feldhaar, H. (2012) Insects, 3, 553–572, Endosymbiont tolerance and control within insect hosts.

102. Ghildiyal, M. and Zamore, P. D. (2009) Nat Rev Genet, 10, 94–108, Small silencing RNAs: an expanding universe.

103. Hammond, S. M., Bernstein, E., Beach, D., et al. (2000) Nature, 404, 293–296, An RNA-directed nuclease mediates post-transcriptional gene silencing in Drosophila cells.

104. Pal-Bhadra, M., Bhadra, U. and Birchler, J. A. (2002) Mol Cell, 9, 315–327, RNAi related mechanisms affect both transcriptional and posttranscriptional transgene silencing in Drosophila.

105. Liu, Q., Rand, T. A., Kalidas, S., et al. (2003) Science (80-.)., 301, 1921–1925, R2D2, a bridge between the initiation and effector steps of the Drosophila RNAi pathway.

106. Aravin, A. A., Klenov, M. S., Vagin, V. V, et al. (2004) Mol Cell Biol, 24, 6742–6750, Dissection of a natural RNA silencing process in the Drosophila melanogaster germ line.

107. Lee, Y. S., Nakahara, K., Pham, J. W., et al. (2004) Cell, 117, 69–81, Distinct roles for Drosophila Dicer-1 and Dicer-2 in the siRNA/miRNA silencing pathways.

108. Okamura, K., Ishizuka, A., Siomi, H., et al. (2004) Genes Dev, 18, 1655–1666, Distinct roles for Argonaute proteins in small RNA-directed RNA cleavage pathways.

109. Saito, K., Ishizuka, A., Siomi, H., et al. (2005) PLoS Biol, 3, e235, Processing of pre-microRNAs by the Dicer-1-Loquacious complex in Drosophila cells.

110. Scott, J. G., Michel, K., Bartholomay, L., et al. (2013) J. Insect Physiol., Towards the elements of successful insect RNAi.

111. Christiaens, O. and Smagghe, G. (2014) Curr. Opin. Insect Sci., 6, 15–21, The challenge of RNAi-mediated control of hemipterans.

112. Kola, V. S. R., Renuka, P., Madhav, M. S., et al. (2015) Front. Physiol., 6, 119, Key enzymes and proteins of crop insects as candidate for RNAi based gene silencing.

113. Bernstein, E., Caudy, A. A., Hammond, S. M., et al. (2001) Nature, 409, 363–366, Role for a bidentate ribonuclease in the initiation step of RNA interference.

114. Knight, S. W. and Bass, B. L. (2001) Science (80-.)., 293, 2269–2271, A role for the RNase III enzyme DCR-1 in RNA interference and germ line development in Caenorhabditis elegans.

115. Lee, Y., Ahn, C., Han, J., et al. (2003) Nature, 425, 415–419, The nuclear RNase III Drosha initiates microRNA processing.

116. Leuschner, P. J., Obernosterer, G. and Martinez, J. (2005) Curr Biol, 15, R603–5, MicroRNAs: Loquacious speaks out.

117. Yeom, K. H., Lee, Y., Han, J., et al. (2006) Nucleic Acids Res, 34, 4622–4629, Characterization of DGCR8/Pasha, the essential cofactor for Drosha in primary miRNA processing.

118. Liu, X., Jiang, F., Kalidas, S., et al. (2006) RNA, 12, 1514–1520, Dicer-2 and R2D2 coordinately bind siRNA to promote assembly of the siRISC complexes.

119. Liu, X., Park, J. K., Jiang, F., et al. (2007) RNA, 13, 2324–2329, Dicer-1, but not Loquacious, is critical for assembly of miRNA-induced silencing complexes.

120. Okamura, K., Robine, N., Liu, Y., et al. (2011) Mol Cell Biol, 31, 884–896, R2D2 organizes small regulatory RNA pathways in Drosophila.

121. Taning, C. N. T., Andrade, E. C., Hunter, W. B., et al. (2016) Sci. Rep., 6, 38082, Asian Citrus Psyllid RNAi Pathway – RNAi evidence.

122. Czech, B., Malone, C. D., Zhou, R., et al. (2008) Nature, 453, 798–802, An endogenous small interfering RNA pathway in Drosophila.

123. Haac, M. E., Anderson, M. A., Eggleston, H., et al. (2015) Nucleic Acids Res, 43, 3688–3700, The hub protein loquacious connects the microRNA and short interfering RNA pathways in mosquitoes.

124. Hammond, S. M., Boettcher, S., Caudy, A. A., et al. (2001) Science (80-.)., 293, 1146–1150, Argonaute2, a link between genetic and biochemical analyses of RNAi.

125. Liu, J., Carmell, M. A., Rivas, F. V, et al. (2004) Science (80-.)., 305, 1437–1441, Argonaute2 is the catalytic engine of mammalian RNAi.

126. Meister, G. and Tuschl, T. (2004) Nature, 431, 343–349, Mechanisms of gene silencing by double-stranded RNA.

127. Verdel, A., Jia, S., Gerber, S., et al. (2004) Science (80-.)., 303, 672–676, RNAi-mediated targeting of heterochromatin by the RITS complex.

128. Kennerdell, J. R., Yamaguchi, S. and Carthew, R. W. (2002) Genes Dev, 16, 1884–1889, RNAi is activated during Drosophila oocyte maturation in a manner dependent on aubergine and spindle-E.

129. Jaubert-Possamai, S., Rispe, C., Tanguy, S., et al. (2010) Mol Biol Evol, 27, 979–987, Expansion of the miRNA pathway in the hemipteran insect Acyrthosiphon pisum.

130. Zhang, Y., Wang, Y., Wang, L., et al. (2016) Pestic. Biochem. Physiol., 127, 21–27, Knockdown of NADPH-cytochrome P450 reductase results in reduced resistance to buprofezin in the small brown planthopper, Laodelphax striatellus (fall{e}n).

131. McLean, K. J., Luciakova, D., Belcher, J., et al. In Monooxygenase, Peroxidase and Peroxygenase Properties and Mechanisms of Cytochrome P450; Springer, 2015; pp. 299–317.

132. Feyereisen, R. (2006) Biochem. Soc. Trans., 34, 1252–1255, Evolution of insect P450.

133. Schama, R., Pedrini, N., Juarez, M. P., et al. (2016) Insect Biochem. Mol. Biol., 69, 91–104, Rhodnius prolixus supergene families of enzymes potentially associated with insecticide resistance.

134. Richards, S., Gibbs, R. A., Weinstock, G. M., et al. (2008) Nature, 452, 949–55, The genome of the model beetle and pest Tribolium castaneum.

135. Saha, S., Hunter, W., Mueller, L., et al. Diaphorina citri genome assembly Diaci 1.9 2017.

136. Chaisson, M. J. and Tesler, G. (2012) BMC Bioinformatics, 13, 238, Mapping single molecule sequencing reads using Basic Local Alignment with Successive Refinement (BLASR): Theory and Application.

137. Stanke, M., Steinkamp, R., Waack, S., et al. (2004) Nucleic Acids Res., 32, W309--W312, AUGUSTUS: a web server for gene finding in eukaryotes.

138. Slater, G. S. C. and Birney, E. (2005) BMC Bioinformatics, 6, 31, Automated generation of heuristics for biological sequence comparison.

139. Kim, D., Pertea, G., Trapnell, C., et al. (2013) Genome Biol., 14, R36, TopHat2: accurate alignment of transcriptomes in the presence of insertions, deletions and gene fusions.

140. Trapnell, C., Roberts, A., Goff, L., et al. (2012) Nat. Protoc., 7, 562–578, Differential gene and transcript expression analysis of RNA-seq experiments with TopHat and Cufflinks.

141. Haas, B. J., Papanicolaou, A., Yassour, M., et al. (2013) Nat. Protoc., 8, 1494–1512, De novo transcript sequence reconstruction from RNA-seq using the Trinity platform for reference generation and analysis.

142. Jones, P., Binns, D., Chang, H.-Y., et al. (2014) Bioinformatics, 30, 1236–1240, InterProScan 5: genome-scale protein function classification.

143. Wu, T. D. and Watanabe, C. K. (2005) Bioinformatics, 21, 1859–1875, GMAP: a genomic mapping and alignment program for mRNA and EST sequences.

144. Kim, D., Langmead, B. and Salzberg, S. L. (2015) Nat. Methods, 357–360, HISAT: a fast spliced aligner with low memory requirements.

145. Pertea, M., Pertea, G. M., Antonescu, C. M., et al. (2015) Nat. Biotechnol., 290–295, StringTie enables improved reconstruction of a transcriptome from RNA-seq reads.

